# Psoriasis-like inflammation induces structural and functional changes in mitochondria

**DOI:** 10.1101/2024.09.05.611357

**Authors:** Gabrielė Kulkovienė, Martyna Uldukytė, Sofiya Haluts, Milda Kairytė, Jonas Šoliūnas, Viktorija Šalčiūtė, Rūta Inčiūraitė, Jurgita Skiecevičienė, Monika Iešmantaitė, Ramunė Morkūnienė, Aistė Jekabsone

## Abstract

Mitochondrial structural and functional changes accompany psoriasis, yet the mitochondrial response to psoriatic inflammation in keratinocytes and fibroblasts remains unexplored. In this study, we investigated the effect of psoriasis-like inflammation (PLI) induced by a cytokine cocktail (interleukin (IL)-17A, IL-22 and tumour necrosis factor (TNF)-α) on mitochondrial network morphology and function in cultured keratinocytes (HaCaT) and fibroblasts (BJ-5ta). In both cell types, PLI triggered expression of psoriasis-related Elafin and high amounts of cytokines (IL-1, IL-6), interferons (IFN-α, IFN-β, IFN-γ) and chemokines (C-C motif chemokine 5 (CCL5) and IL-8), accompanied by increased mitochondrial membrane potential, reactive oxygen species (ROS) production, respiration suppression, network fragmentation, swelling, and cristae disassembly. Stimulated emission depletion (STED) nanoscopy revealed the disappearance of mitochondrial cristae in response to PLI, with the process starting more quickly and being more pronounced in keratinocytes than in fibroblasts. These findings highlight cell-specific mitochondrial responses to psoriatic inflammation, guiding future investigations towards new pharmacological targets for managing psoriasis.

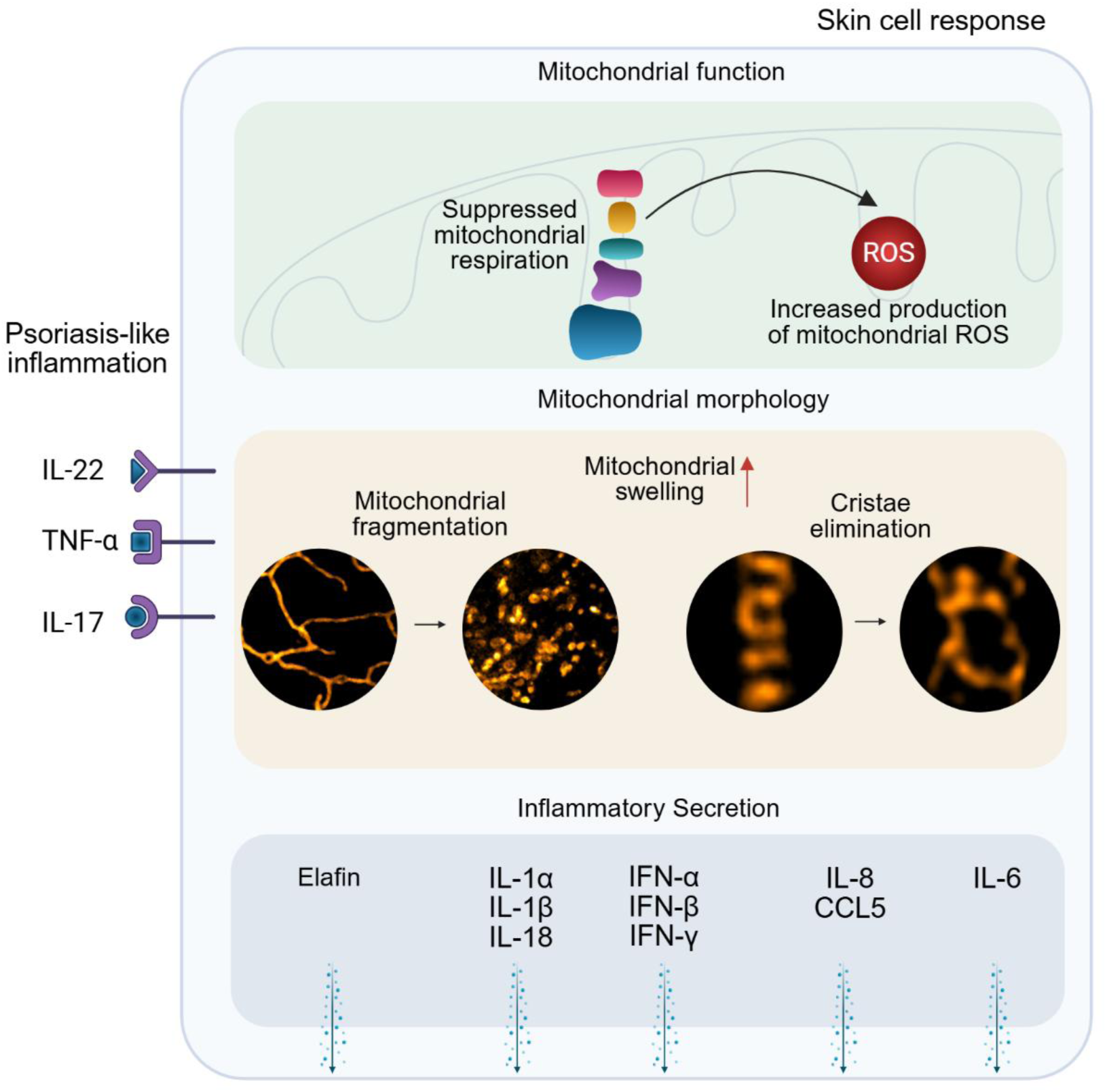

## Introduction

Psoriasis is a chronic recurrent inflammatory skin disease characterised by immune system dysregulation and epidermal hyperproliferation. During psoriasis, immune cell-derived proinflammatory cytokines affect keratinocytes and fibroblasts, resulting in psoriatic inflammation, which promotes the formation of lesions [1]. A cytokine cocktail comprising tumour necrosis factor (TNF)-α, interleukin (IL)-17A, and IL-22 can mimic psoriasis-like inflammation (PLI) *in vitro* [2,3]. PLI increases the production of antimicrobial peptides (Elafin, β-defensin, LL-37), which act as potential autoantigens for immune cells at early disease stage [4]. For instance, *PI3*, encoding Elafin, is not expressed in the normal human epidermis [5], and the secretion of this protein is linked with disease severity [6].

Systemic biological agents targeting psoriasis-related cytokines are considered the most advanced treatment option for psoriasis. Despite their efficacy, biological therapies often exhibit significant variability in individual responses, come with high costs and carry the risk of severe side effects [7]. A better understanding of the disease pathogenesis would enable new, more effective treatment strategies.

Recent evidence suggests that mitochondrial involvement is a key factor in psoriasis development, with mitochondrial reactive oxygen species (MitoROS) and network organisation identified as significant contributors to the disease process. *In vivo* studies revealed MitoROS-dependent dendritic cell activation, while MitoROS scavenging effectively reduced epidermal hyperplasia and the infiltration of inflammatory cells [8]. Mitochondrial fragmentation in macrophages induced psoriasis-related cytokines, facilitating their movement to the epidermis [9]. However, it remains unclear if such mitochondrial transformation occurs in keratinocytes and fibroblasts, the critical players in psoriasis pathogenesis. Understanding the role of MitoROS and mitochondrial network-controlling factors in these cells could offer potential therapeutic targets for fighting psoriatic inflammation.

Mitochondria respond to stress by changing their structure, with cristae architecture playing a particular role in this process [10]. Recent advances in super-resolution microscopy enabled the visualisation of cristae dynamics in living cells and can provide new data about the role of mitochondrial structural changes in psoriasis.

In our study, we investigated and compared the mitochondrial response to PLI in fibroblasts and keratinocytes, focusing on mitochondrial function, network, and cristae architecture. PLI was induced using a cytokine cocktail TNF-α, IL-17A, and IL-22, with model validation achieved through Elafin mRNA expression and the secretion of psoriasis-related cytokines. We assessed mitochondrial network organisation and cristae structure using STED nanoscopy, alongside measurements of mitochondrial membrane potential, ROS production and functional changes.

## Results

### Induction and validation of psoriasis-like inflammation in keratinocytes and fibroblasts

After treating human skin keratinocytes (KCs) and fibroblasts (FCs) with IL-17, IL-22, and TNF-α for 24 hours to induce PLI, the metabolic activity of KCs declined by 11.5% and that of FCs - by 31% compared to the untreated control. After this decline, the metabolic activity of the cells remained stable for 48 and 72 hours (Figure 1a). Cell viability assessment by Hoechst 33342 and propidium iodide double nuclear staining showed no significant changes in the viability of the cells (Figure 1c). However, the total cell numbers were lower after PLI treatment than in the untreated controls. The numbers of PL-KCs were 34% smaller after 48-hour and 36% - after 72-hour PLI treatment, and those of PL-FCs decreased by 28% after 48 hours with PLI and returned to baseline levels after 72 hours (Figure 1b). These results indicate that PLI does not cause cell death but induces a sustained decline in cell proliferation over time. As such, this model remains robust enough to be suitable for further experimental investigation.

**Figure 1.**
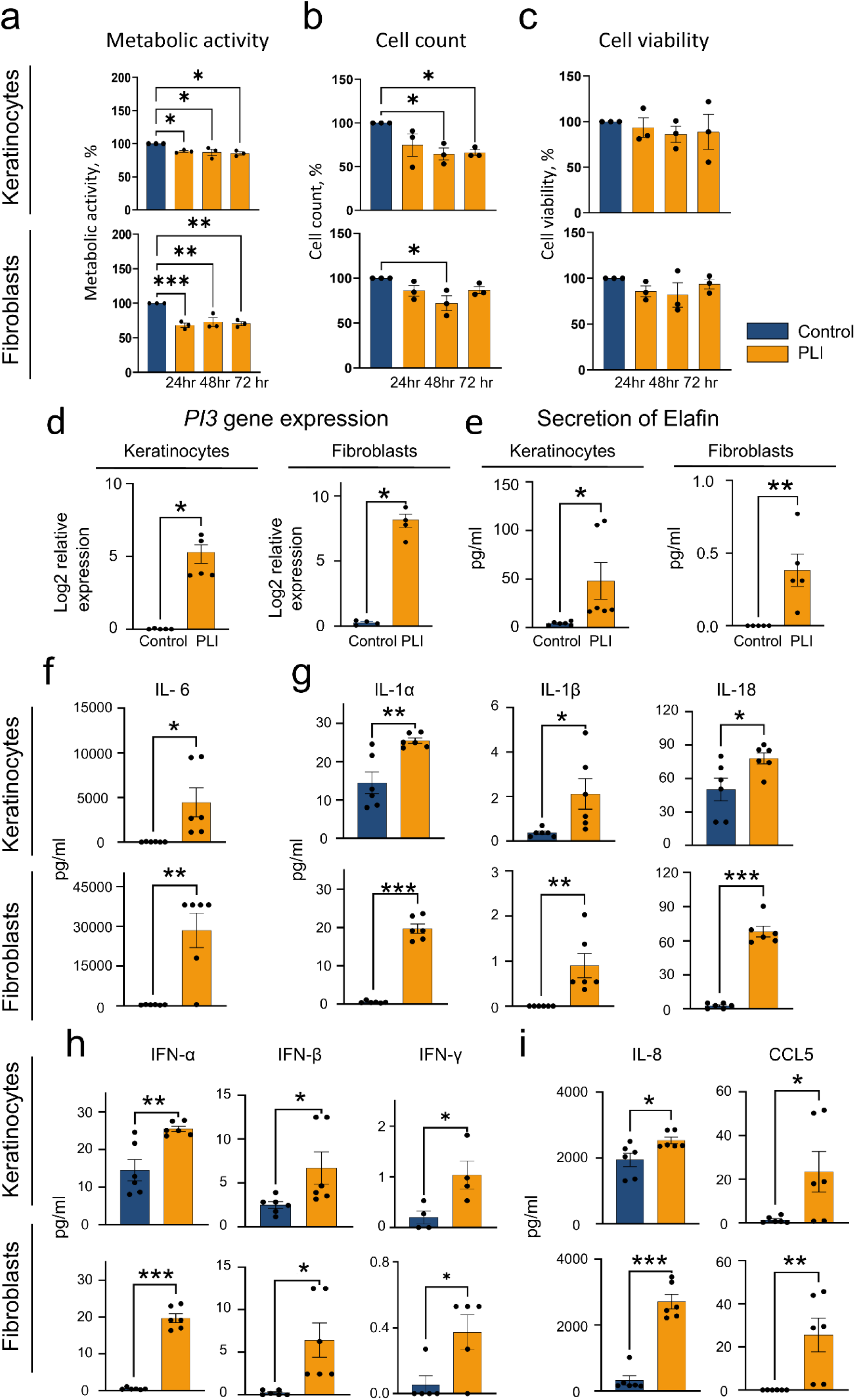
Induction and validation of psoriasis-like inflammation (PLI) in keratinocytes and fibroblasts. Metabolic activity (a), cell number (b) and cell viability (c) change over time after treatment with PLI-inducing cytokines. Cell viability was expressed as the ratio of live and necrotic cells in a population. Data: mean± SEM. Statistics: One-way ANOVA Dunnett’s test, n=3.(d) *PI3* gene expression assessed by RT-qPCR; n=5 for keratinocytes (KCs), n=4 for fibroblasts (FCs). Fold change presented in log_2_ scale. (e) Secretion of Elafin. n=6 for KCs, n=5 for FCs. (f) Secretion of IL-6. (g) Secretion of IL-1 family members. (h) Secretion of interferons IFN-α, IFN-β, IFN-γ. (i) Secretion of chemokines IL-8 and CCL5. Protein levels were quantified by Luminex multiplex assay. Data: mean±SEM. Statistics: unpaired, two-tailed Student’s t-test. n=6 for all analytes, except for IFN-γ (n=4 for KCs, n=5 for FCs). \**p* < 0.05, ** p <0.01, *** p<0.001.

Next, psoriatic phenotype induction in the PLI-treated cells was assessed by measuring the expression and secretion of psoriatic markers. PLI treatment significantly upregulated the psoriatic skin marker PI3 expression, with a 5.27-fold increase in KCs and a 7.8-fold increase in FCs (log2, Figure 1d). Moreover, Elafin secretion was markedly elevated, reaching 47.98 ±18.93 pg/ml in KCs and 0.38±0.11 pg/ml in FCs (Figure 1e). In untreated controls, protein expression and secretion were near zero or undetectable. The observed increase in Elafin confirms the PLI-induced changes in KCs and FCs; therefore, we refer to them as psoriatic-like (PL) cells in this study.

Psoriasis indicating marker assessment was followed by testing levels of disease-propagating cytokines. Under PLI conditions, both FCs and KCs secreted significantly higher amounts of proinflammatory mediators than untreated controls. These included IL-6 (Figure 1f) and cytokines from the IL-1 family (IL-1α, IL-1β, IL-18, Figure 1g). Among these, IL-6 - a key mediator of Th17-driven cutaneous inflammation essential for psoriasis initiation – showed the most substantial increase in both cell types, with notably higher levels in PL-FCs than PL-KCs (Figure 1f). Similarly, IL-1α concentrations increased in both cell types under PLI, with more pronounced levels in PL-KCs (Figure 1g).

Inflammasome-related cytokines IL-18 and IL-1β also rose after PLI induction, with IL-1β being significantly elevated in PL-KCs, while IL-18 showed higher levels in PL-FCs (Figure 1g). Among interferons, IFN-α displayed the most substantial increase in both cell types, followed by moderate rises in IFN-β and minimal changes in IFN-γ (Figure 1h).

Chemokines CCL5 and IL-8 were highly expressed in PL cells, with IL-8 levels being higher in PL-FCs than in PL-KCs. Meanwhile, CCL5 concentrations increased similarly across both cell types (Figure 1i).

While both cell types secreted higher amounts of cytokines and chemokines after PLI induction, a notable distinction emerged in their secretion profiles. PL-FCs produced higher levels of IL-6 and IL-8 compared to PL-KCs, while PL-KCs secreted more IL-1β than PL-FCs. The results suggest that both cell types participate in PLI propagation with distinct roles.

### Psoriasis-like inflammation disrupts mitochondrial network

Aiming to test PLI effect on mitochondrial network integrity in KCs and FCs, we performed mitochondrial morphometry analysis using Mito Live Orange dye, STED nanoscopy and MiNa toolset in Fiji (Figure 2a). As a positive assay control, we used the mitochondrial stressor FCCP, which makes the mitochondrial inner membrane proton-permeable, leading to membrane potential loss and mitochondrial fragmentation [11–13].

**Figure 2.**
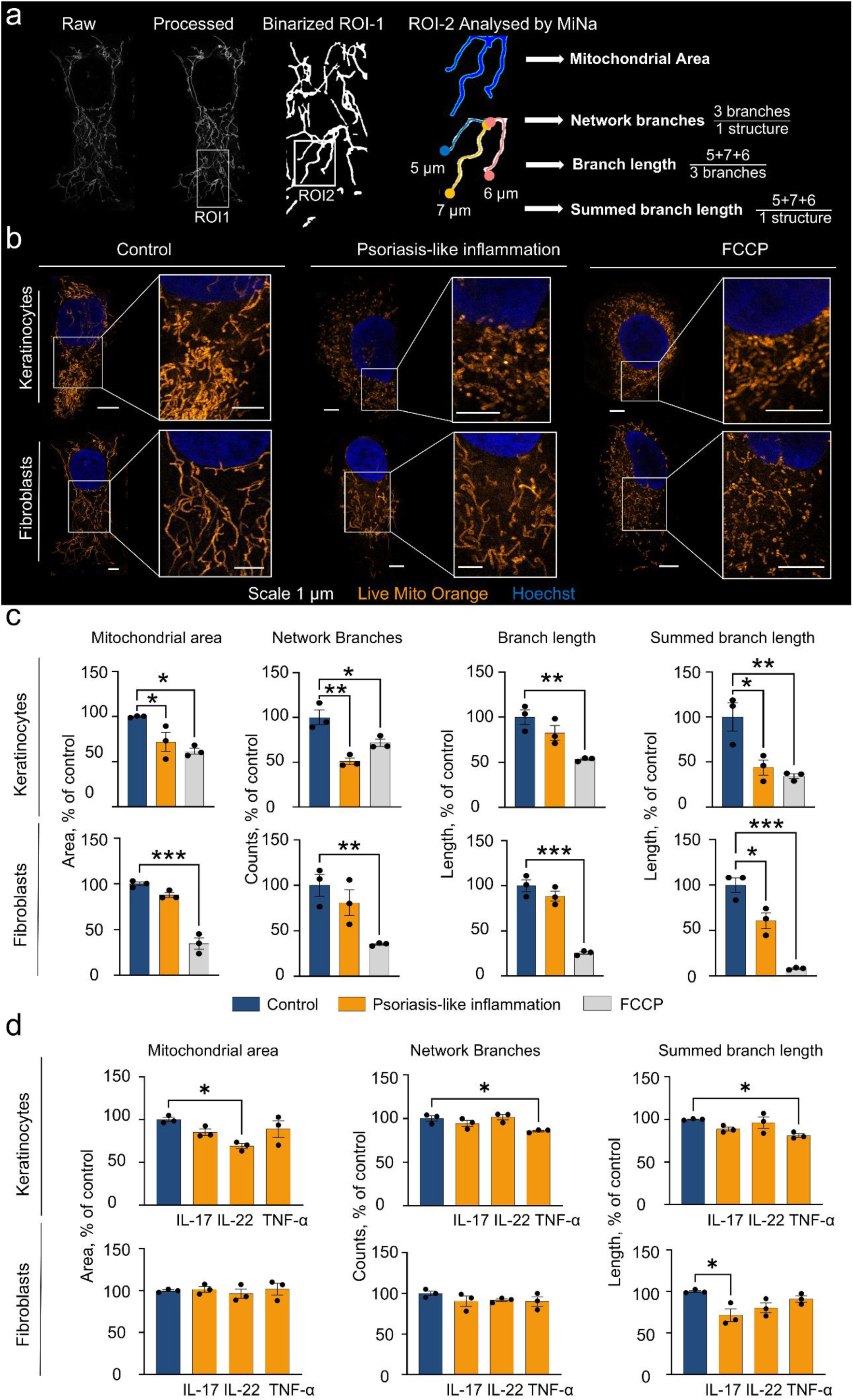
Mitochondrial morphometry of keratinocytes and fibroblasts after 24-hour treatment with psoriasis-like inflammation (PLI)-inducing cytokines. (a) Steps of mitochondrial network analysis and explanation of the calculated parameters. Images were processed using unsharp masking, CLAHE, and median filtering techniques. MiNa plugin in Fiji binarised the images and calculated the mitochondrial area. Afterwards, it skeletonised binary images and calculated the mean of network branches, mean branch length, and summed branch length. ROI – Region of interest. (b) Representative STED nanoscopy images of cells labelled with Live Mito Orange. FCCP was used as a positive control. Scale-1µm. n=3 (c) Quantitative results of mitochondrial network parameters. (d) Mitochondrial morphometry of keratinocytes and fibroblasts treated with individual IL-17, IL-22 and TNF-α for 24 hours. Data: mean±SEM. Statistics: One-way ANOVA Dunnett’s test. n=3. *p< 0.05, ** p <0.01, ***p<0.001.

Representative images (Figure 2b) show a dramatic fragmentation in PL-KCs, where mitochondria appear vesicle-shaped rather than network-like. At the same time, PL-FCs maintained the network shape but showed shorter structures. Quantitative analysis revealed a significant decrease in the mitochondrial area (28%) and network branches (48%) in PL-KCs compared to control KCs (Figure 2c), while in PL-FCs, these parameters remained unchanged. Both cell types displayed a 56% and 39% decrease in summed branch length in PL-KCs and PL-FCs, respectively, while overall branch length remained unchanged. These findings indicate an increased number of isolated mitochondrial networks, suggesting fragmentation in both PL-cell types.

We compared PLI-induced changes to FCCP-induced fragmentation as a reference for extreme mitochondrial stress (Figure 2c). In KCs, FCCP caused a 38% reduction in mitochondrial area, 28% in network branches, 46% in branch length and 65% in summed branch length. PLI-induced changes were similar but showed a more substantial reduction in network branches (48% vs FCCP’s 28%), indicating significant mitochondrial fragmentation in PL-KCs. In FCs, FCCP caused a decrease of 65% in mitochondrial area, 64% in network branches, 74% in branch length, and 91% in summed branch length. These changes were more pronounced than those induced by PLI, indicating relatively moderate mitochondrial fragmentation in PL-FCs.

To determine individual cytokine’s role in PLI-induced mitochondrial changes, we exposed cells to either IL-17A, IL-22 or TNF-α (Figure 2d). IL-22 and TNF-α affected network parameters in KCs, while IL-17A primarily affected FCs, inducing moderate mitochondrial fragmentation. Notably, the combination of three cytokines caused more pronounced changes than each cytokine individually.

The data indicate that PLI-inducing cytokines synergise at mitochondria of both cell types, with KCs exhibiting significantly more pronounced network fragmentation than FCs.

### Psoriasis-like inflammation induces mitochondrial swelling and affects cristae architecture

Mitochondrial network changes suggest a possible impact of PLI on mitochondrial cristae organisation; thus, we tested if PLI alters cristae morphology and compared changes in PL-KCs and PL-FCs, using Live Mito Orange and STED nanoscopy for super-resolution imaging. Cristae area, density and spacing were assessed following steps depicted in Figure 3a.

**Figure 3.**
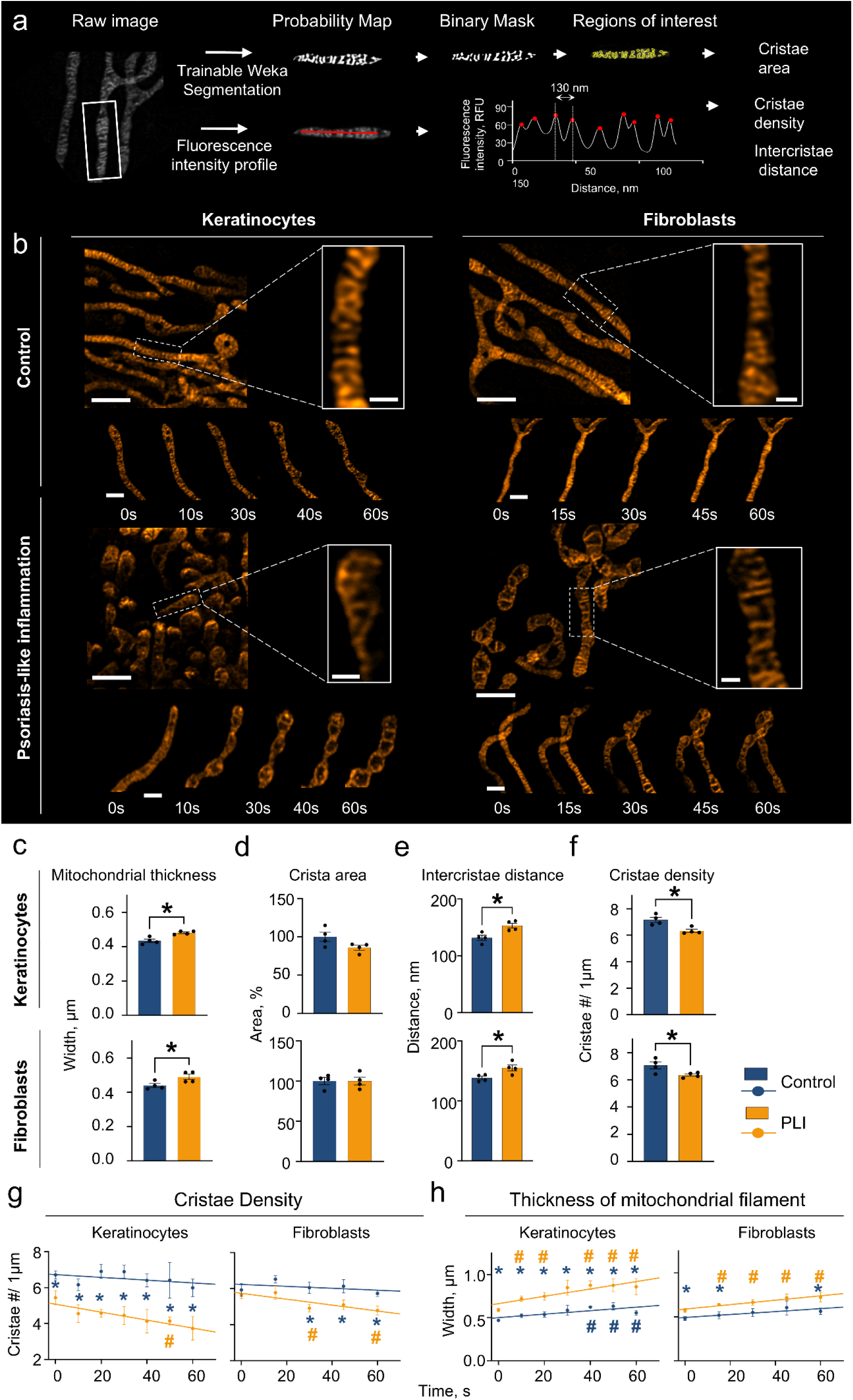
Mitochondrial cristae characteristics of keratinocytes and fibroblasts treated psoriasis-like inflammation (PLI)-inducing cytokines for 24 hours. (a) – Cristae analysis scheme. Cristae area was evaluated using Trainable Weka Segmentation, spacing - as average peak (cristae) distances along 1-2µm fluorescence intensity profiles and density as peaks/1µm. (b) – Overview and time-lapse (Scales: 1µm, 200nm). (c) – Mitochondrial filament thickness. (d) – Average crista area. (Control = 100%). (e) – Intercristae distance. (f) – Cristae density. (g) – Cristae density and (h) – mitochondrial thickness change over 60s. Data: mean ± SEM. Statistics: unpaired, two-tailed Student’s t-test. #,*p< 0.05, n=4. In (g), (h): blue asterisk (*) indicates a significant difference vs. control at the same time point. A dash (#) - a comparison with the initial time point (0s) within the control (blue #) and the PLI (yellow #) groups.

Under PLI, mitochondria appeared swollen with sparsely arranged cristae in both cell types compared to controls (Figure 3b, upper part). Quantitative analysis confirmed a significantly increased mitochondrial width by 10.9% in PL-KCs and 11.4% in PL-FCs (Figure 3c); without changes in cristae area (Figure 3d). Intercristae distance increased by 16.2% in PL-KCs and 12.2% in PL-FCs (Figure 3e). Lastly, the cristae elimination process was confirmed by evaluating the cristae count per 1µm (cristae density), which decreased by 11.5% in PL-KCs and 10.1% in PL-FCs (Figure 3f).

To determine whether mitochondrial swelling was finite or continuous, we measured changes over 60 seconds. Time-lapse images (Figure 3b, lower part) showed ongoing cristae reduction and mitochondrial filament swelling in PL cells. In PL-KCs, mitochondrial filaments expanded by 20-55% between 10-60s (Figure 3h). Control KCs showed less swelling (18-34%) towards the end of the recording (40-60s). In PL-FCs, mitochondrial filaments expanded by 11-26% between 15-60s, with no further swelling in control FCs. Cristae density in PL cells gradually decreased (Figure 3g): by 23% in PL-KCs at 50s and by 13-15% in PL-FCs between 30-60s. Control KCs and FCs showed no change. Interestingly, cristae density was significantly lower at the moment of observation (0s) in PL-KCs compared to controls, while no initial difference was observed in PL-FCs vs. control. However, cristae density in PL-FCs significantly decreased after 30s, suggesting a delayed process compared to PL-KCs.

Summarising, PLI caused persistent mitochondrial swelling and a gradual decrease in cristae density in PL cells, with cristae elimination evident early in PL-KCs and delayed in PL-FCs.

### Psoriasis-like inflammation increases mitochondrial membrane potential, ROS production and alters bioenergetic activity in keratinocytes and fibroblasts

Mitochondrial network changes are tightly associated with shifts in mitochondrial respiration and ROS production. Given our previous observation that mitochondrial structural changes progressed faster in PL-KCs while delayed in PL-FCs, we aimed to assess whether this correlated with functional alterations over time. Therefore, we further determined mitochondrial membrane potential (Δψ_m_) using Tetramethylrhodamine methyl ester (TMRM) (Figure. 4a, c), MitoROS production – with MitoSOX (Figure 4b, d, e, f) and examined bioenergetic activity using Seahorse XFp MitoStress (Figure 5a, b) assay after prolonged PLI exposure.

**Figure 4.**
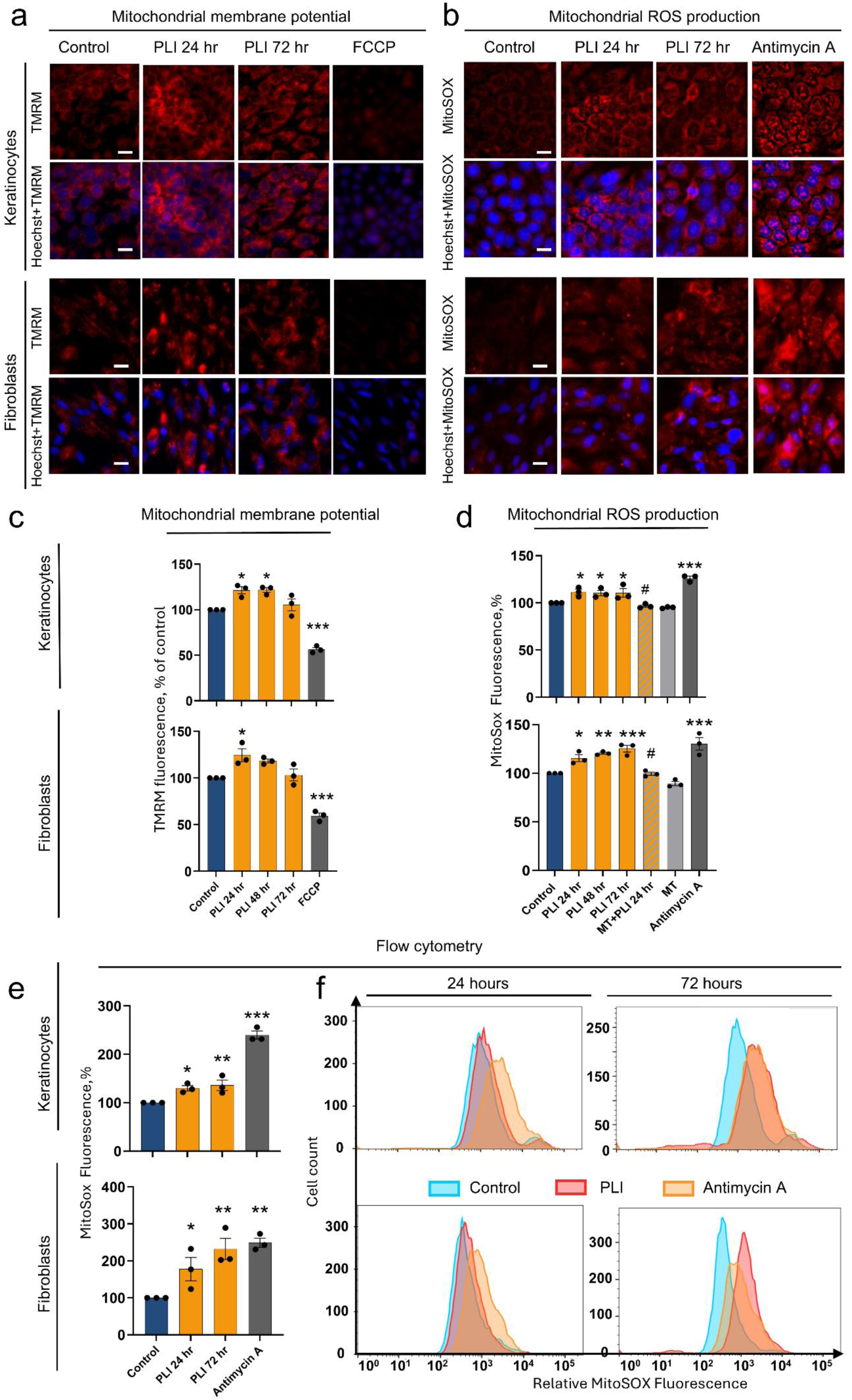
Mitochondrial membrane potential and reactive oxygen species (ROS) production in psoriasis-like inflammation (PLI) induced keratinocytes and fibroblasts over time. (a) Representative fluorescent microscopy images of cells labelled with TMRM, with FCCP – used as a control to induce mitochondrial membrane depolarisation. (b) Representative fluorescent microscopy images of cells labelled with MitoSOX, with Antimycin A as a control for increased ROS production. Scale in (a) and (b) - 20µm. (c) Quantitative TMRM results. (d) Quantitative results of MitoSOX, evaluated by fluorescent microscopy. MitoTempo (MT) was used as a negative control. (e) Quantitative results of MitoSOX, evaluated by flow cytometry. (f) Representative histograms of MitoSOX fluorescence. Data: mean ± SEM. Statistics: in (d) One-way ANOVA Fisher’s LSD, in (c), (e) - Dunnett’s test. An asterisk (*) indicates a significant difference vs control, while a dash (#) indicates a significant difference vs PLI 24 hours. n=3. *p < 0.05, ** p <0.01, *** p<0.001.

**Figure 5.**
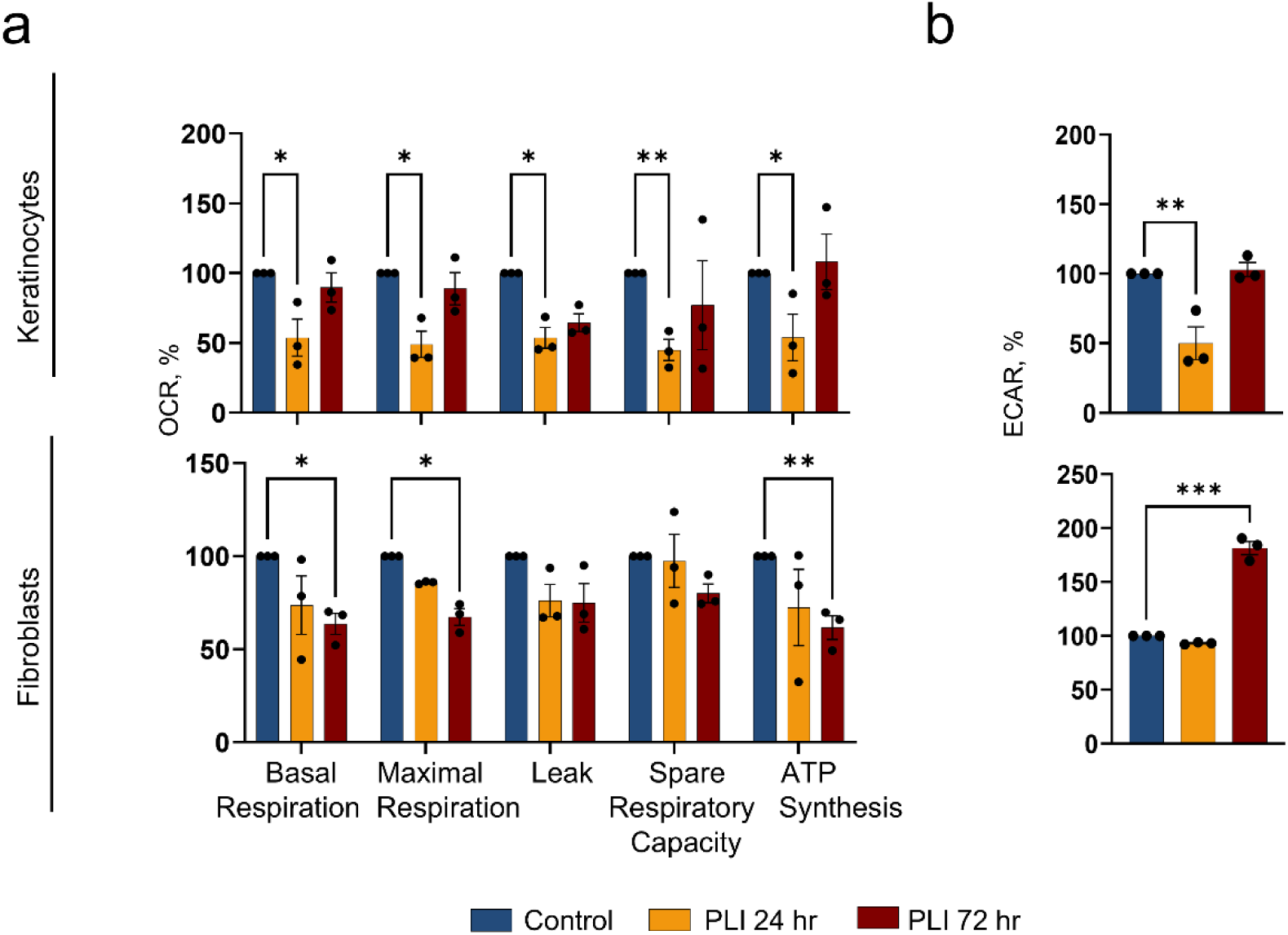
Bioenergetic activity in psoriasis-like inflammation (PLI) induced keratinocytes and fibroblasts over time. (a) Summarised oxygen consumption rate (OCR) parameters. (b) Extracellular acidification rate (ECAR) represents glycolysis efficiency. Data: mean ± SEM Statistics: Two-way ANOVA, Dunnett’s test. n=3. *p< 0.05, ** p <0.01, ***p<0.001.

After 24 hours of PLI induction, Δψ_m_ increased by 24% in PL-FCs and 21% in PL-KCs, where it remained at a similar level after 48 hours. By 72 hours, both cell lines gradually returned to baseline (Figure 4a, c). Unlike this transient Δψ_m_ increase, MitoROS production demonstrated a progressive increase in PL-FCs, rising by 15% at 24 hours, 20% at 48 hours and 25% at 72 hours (Figure 4d). In contrast, MitoROS in PL-KCs peaked at 24 hours (12.5%), stabilising around 11% at 48 and 72 hours (Figure 4d). MitoROS measurement by flow cytometry corroborated the fluorescent microscopy results while revealing more pronounced differences between groups (Figure 4e, f). In FCs, PLI induced MitoROS production by 77% after 24 hours and 131% after 72 hours. In contrast, PL-KCs showed a more modest response, with MitoROS increasing by 25% after 24 and 39% after 72 hours (Figure 4e).

Changes in mitochondrial respiration and glycolysis efficiency were detected at different time points in PL-KCs and PL-FCs. PL-KCs exhibited a significant decrease in overall mitochondrial oxygen consumption rate (OCR) after 24-hour PLI induction. Basal respiration decreased by 46%, leading to a notable drop in ATP production, while maximal respiration was suppressed by 50% (Fig. 5a). Similarly, spare respiratory capacity in PL-KCs was 55% lower than in control. The glycolytic activity was also suppressed in PL-KCs, dropping to 50% of the control cells (Figure 5b), as confirmed by the XFp Glycolysis Stress Test (data not shown). These effects diminished by 72 hours. In contrast, in PL-FCs, mitochondrial respiration and glycolysis efficiency remained unaltered after 24 hours of PLI exposure. However, after 72 hours, mitochondrial functions have declined, with reductions of 36% in basal, 32% in maximal respiration, and a 38% drop in ATP production. Moreover, this mitochondrial suppression was accompanied by a glycolytic efficiency increase of 80%.

Our findings suggest that in PL-KCs, the response is immediate, with a significant rise in MitoROS production and a decrease in mitochondrial respiration and glycolysis within 24 hours. Conversely, in FCs, PLI triggers a delayed response. MitoROS increase, likely driven by the rise in Δψ_m_, progressively accumulates over 72 hours, eventually impacting mitochondrial respiration and glycolysis efficiency.

### MitoTempo and Mdivi1 effect on Drp1 activation and psoriasis-like inflammation induced mitochondrial fragmentation

Dynamin-related protein 1 (Drp1) regulates mitochondrial fission by translocating to the outer mitochondrial membrane, oligomerising and undergoing GTPase-driven constriction, which results in mitochondrial division. Drp1 activation involves post-translational modifications, notably the phosphorylation of Ser616. We performed a Western Blot analysis to verify whether PLI-induced fragmentation is Drp1-dependent.

PLI significantly induced both Drp1 and its phosphorylation at Ser616 (pDrp1) proportionally in KCs (Fig. 6a, quantitative data not shown) without changing their ratio. This suggests that PLI has a direct impact on Drp1 expression and activation, leading to increased mitochondrial fission activity. Mdivi1, a known Drp1 inhibitor, lowered the pDrp1/Drp1 ratio (Fig. 6a, c) in PL-KCs while restoring mitochondrial integrity, as evidenced by normalised summed branch length (Fig. 6d, e). These results indicate that the fragmentation induced by PLI is Drp1-dependent. In FCs, neither PLI nor Mdivi1 affected Drp1 phosphorylation (Fig. 6b, c). However, Mdivi1 restored the mitochondrial network and increased the summed branch length by 64% above the control levels in PL-FCs (Fig. 6f).

**Figure 6.**
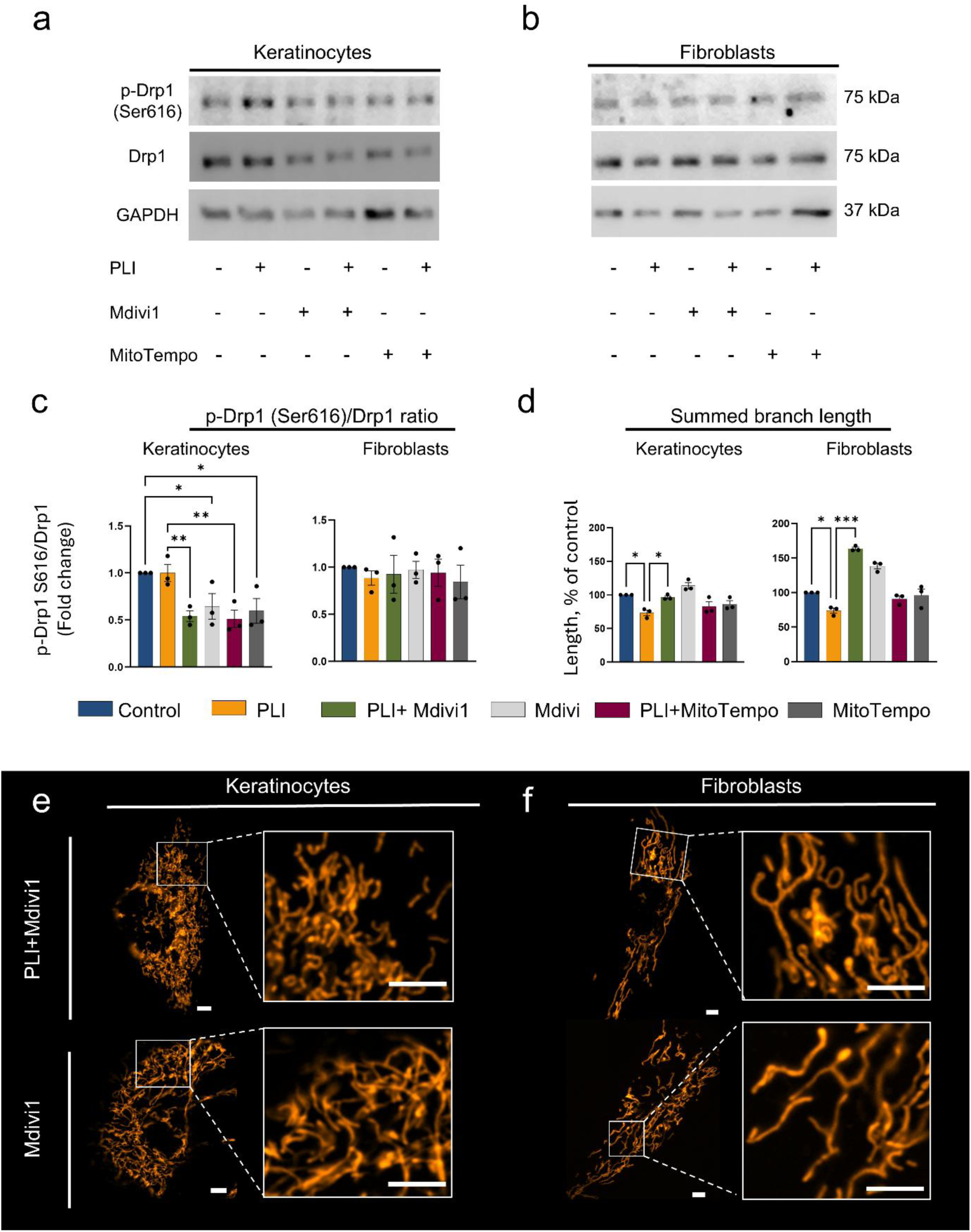
MitoTempo and Mdivi1 effect on Drp1 activation and psoriasis-like inflammation (PLI) induced mitochondrial fragmentation. (a)-(b) Western blots showing protein levels of phosphorylated-Drp1 (Ser616) and total Drp1. (c) pDrp1 (Ser616)/total Drp1 ratio, normalised to loading control GAPDH. (d) Mdivi1 and MitoTempo effect on summed branch length, calculated with MiNa morphology analysis tool in Fiji. (e)-(f) Images representing the restored mitochondrial network in PLI-affected cells after adding the Drp1 inhibitor Mdivi1. Scale - 5µm. Data: mean±SEM. Statistics: One-way ANOVA, Fisher’s LSD test. n=3. *p < 0.05, ** p <0.01, *** p<0.001.

To investigate whether PLI-induced MitoROS contributes to mitochondrial fragmentation, we employed MitoTempo, a MitoROS scavenger. MitoTempo reduced the pDrp1/Drp1 ratio in PL-KCs (Fig. 6a, c) but did not affect PL-FCs (Fig. 6b, c). Mitochondrial morphology remained unchanged in both cell types (Fig. 6d), suggesting that MitoROS contributes to Drp1 activation but is not the sole driver of mitochondrial fragmentation in PL-KCs.

### Increased mitochondrial ROS and fragmentation contribute to the induction of psoriasis-related proinflammatory mediators

Recent studies suggest that releasing inflammatory mediators in immune cells affected by psoriasis depends on MitoROS or mitochondrial fission. To test whether such cytokine release mediation pathways occur in PL-KCs and PL-FCs, we assessed fold changes in the secretion of proinflammatory mediators after treatment with MitoTempo and Mdivi1.

We focused on analytes with high concentrations in PL cell media, specifically IL-6, IL-1α, IL-1β, IL-18, and Elafin. Additionally, we examined secretion changes in PL-cells over 72 hours to test the time-dependent changes in inflammatory mediator secretion (Figure 7a-j).

**Figure 7.**
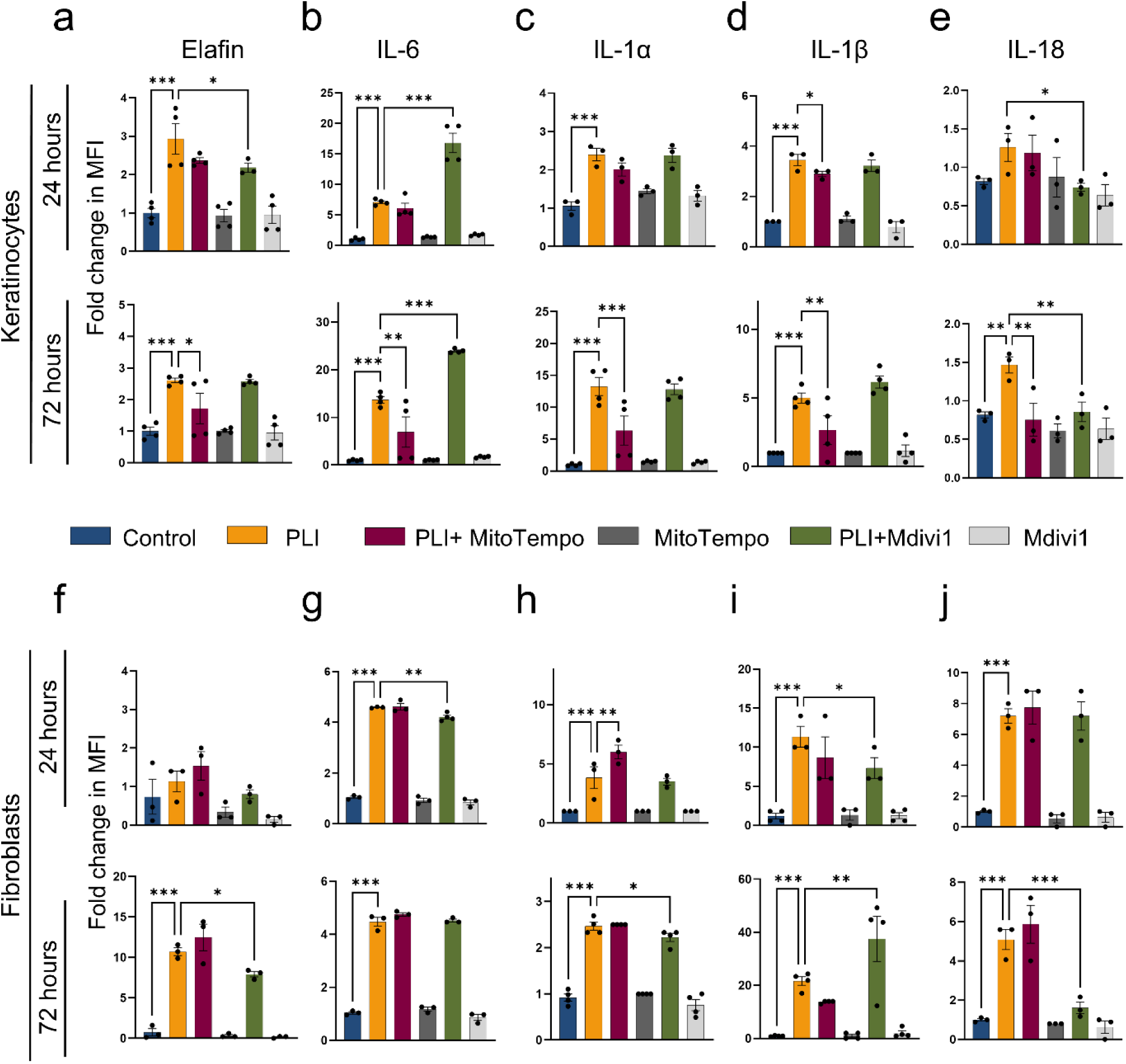
MitoTempo and Mdivi1 effect on proinflammatory mediator secretion in psoriasis-like inflammation (PLI) induced keratinocytes and fibroblasts. Graphs (a)-(e) exhibit proinflammatory mediator secretion in keratinocytes and graphs (f)-(j) – secretion in fibroblasts. Protein levels were quantified by Luminex multiplex assay as mean fluorescent values (MFI); Bar graphs show fold changes in the MFI. Data: mean±SEM. Statistics: One-way ANOVA Fisher’s LSD test. n = 4 for keratinocytes, n = 3 for fibroblasts. *p < 0.05, ** p <0.01, *** p<0.001.

Results showed that the effects of MitoTempo and Mdivi1 on proinflammatory mediators varied by cell type. Notably, MitoTempo significantly reduced the secretion of all mediators in PL-KCs following 72-hour PLI stimulation, specifically lowering Elafin by 0.9-fold (Figure 7a), IL-6 by 6.6-fold (Figure 7b), IL-1α by 6.9-fold (Figure 7c), IL-1β by 2.3-fold (Figure 5d), and IL-18 by 0.6-fold (Figure 7e). However, MitoTempo had no effect on cytokine secretion by PL-FCs. In contrast, Mdivi1 decreased the secretion of most mediators in PL-FCs following 72-hour PLI induction, specifically Elafin by 2.4-fold (Figure 7f), IL-1α by 0.3-fold (Figure 5h) and IL-18 by 3.4-fold (Figure 7j), while IL-6 secretion decreased after 24-hour PLI by 0.38-fold (Figure 7g). In PL-KCs, Mdivi1 showed less impact overall, with two notable exceptions: it reduced Elafin by 0.66-fold following 24-hour PLI (Figure 7a) and IL-18 by 0.93-fold and 1.13-fold following 24- and 72-hour PLI induction, respectively (Figure 7e).

Secretion of some mediators was stimulated by Mdivi1. For instance, it increased IL-6 in PL-KCs by 9.75-fold and 10.41-fold after 24- and 72-hour PLI induction, respectively (Figure 7b). It also raised IL-1β by 13.9-fold in PL-FCs after 72 hours, following an initial reduction of 3.33-fold (Figure 7i). Similarly, MitoTempo elevated IL-1α levels in PL-FCs by 2.16-fold following 24-hour PLI (Figure 7h).

These results reveal that MitoTempo and Mdivi1 can modulate the secretion of proinflammatory mediators, primarily by suppressing their levels, in a manner that varies significantly between different cell types. MitoTempo showed strong inhibitory effects on inflammatory mediator secretion in KCs, while Mdivi1 was more effective in FCs. These findings support the notion that increased MitoROS and fragmented mitochondrial network contribute to the induction of psoriasis-related proinflammatory mediators in a cell type-dependent manner.

## Discussion

In this study, we confirmed that PLI induces cell-type-specific mitochondrial dysfunctions. We found that PLI causes mitochondrial fragmentation, mitochondrial filament swelling, and a gradual decline in cristae density in both cell types, with PL-KCs mitochondria being affected more intensively than those of PL-FCs after 24 hours of PLI induction. Additionally, PLI induces Δψ_m_ and MitoROS production in both cell types. In KCs, these changes are accompanied by an immediate suppression of mitochondrial respiration and glycolysis, whereas in FCs, the response is delayed, with a decrease in mitochondrial respiration and activation of glycolysis. Lastly, our data suggest that the secretion of psoriasis-related proinflammatory mediators may be influenced by increased MitoROS and mitochondrial fragmentation in a cell-type-dependent manner.

Our results suggest that when exposed to psoriatic inflammation, KCs and FCs utilise different mitochondrial adaptations to manage inflammatory stress: KCs respond via the MitoROS pathway, while FCs - through the mitochondrial structure. KCs are targets of external inflammatory stimuli, and their rapid mitochondrial response (including MitoROS increase and respiratory suppression) may be a contributing factor to the progression of inflammatory responses. Previous studies have reported that MitoROS inhibition might serve as a regulatory mechanism. For example, a study [14] showed that IL-17A and IL-22 enhanced IL-1β production by activating NLRP3 inflammasome via ROS in HaCaT cells. We found that, inhibition of MitoROS reduced levels of NLRP3 downstream cytokines, IL-18 and IL-β, in PL-KCs, suggesting a similar mechanism may be involved. In contrast, FCs are better at maintaining mitochondrial integrity and energy homeostasis during inflammation (decreased mitochondrial respiration, increased glycolysis). Mitochondrial fission in PL-FCs may trigger inflammation by regulating proinflammatory mediators. For example, in psoriatic macrophages, fission-related proteins GDAP1L1 and Drp-1 trigger cytokine production through MAPK and NFκB pathways [9], and Drp1 inhibition with Mdivi1 reduces TNF-a-induced NFκB activation [15], alleviating psoriasis-like symptoms [16]. Similarly, our results showed that Drp1 inhibition with Mdivi1 in PL-FCs reduced the secretion of IL-1α, IL-18, and IL-6, suggesting it might also involve NF-κB. While Mdivi1 restored the mitochondrial network, it did not affect Drp1 phosphorylation, suggesting that PL-FCs fragmentation may be regulated by alternative post-translational modifications of the fission machinery. For example, it has been shown that SUMOylation of the major Drp1 receptor, MFF, is necessary to promote mitochondrial fission in mouse embryonic fibroblasts [17]. The effective restoration of the mitochondrial network by Mdivi1 likely results from its inhibition of Drp1’s GTPase activity, preventing fission [18].

While Mdivi1 was shown to effectively inhibit Drp1-mediated mitochondrial fission [16,19–23], it has also been reported to exert off-target effects, including inhibition of mitochondrial complex I, particularly at higher concentrations or with prolonged exposure [24–27]. It was reported that 50 µM Mdivi1 impairs bioenergetics in neurons independently of Drp1 activity, while a lower concentration of 25 µM had no effect on respiratory capacity [24]. The same group also demonstrated that Mdivi1 can directly scavenge free radicals generated by complex I inhibition [26]. In contrast, another study using 100 µM Mdivi1 showed alterations in bioenergetics and increased ROS levels, likely due to reduced Drp1 expression in cardiomyocytes [27]. Furthermore, Marx et al. observed destabilisation of mitochondrial supercomplexes and elevated ROS following either 10 µM Mdivi1 for one week or 50 µM for 24 hours in HeLa and neuronal progenitor cells [25]. These findings suggest that Mdivi1 affects mitochondrial respiratory chain in a dose- and time-dependent manner. Notably, such Mdivi1 interaction with mitochondrial respiratory chain has been observed in neurons, COS-7 cells, cardiomyocytes, and HeLa cells, while evidence from skin cells remains limited. In our study, we applied Mdivi1 at a lower concentration of 25 µM for 24 hours, and this treatment did not affect mitochondrial respiration in KCs and FCs (data not shown). No change in respiration rate suggests that, in our case, complex I inhibition did not occur, and the impact of Mdivi1 on cytokine secretion was most likely related to mitochondrial dynamics. Nonetheless, the possibility of other off-target effects of Mdivi1 cannot be entirely excluded, and future studies, such as Drp1 silencing or gene editing, may help further elucidate the role of this GTPase.

Our results, for the first time, showed that psoriasis-like inflammation causes mitochondrial filament swelling, accompanied by cristae elimination in KCs and FCs. In our previous study, we captured similar changes in a viral infection model of human lung microvascular cells [28], suggesting that this may be a trend of inflammatory response. Transmission electron microscopy (TEM) studies on similar models have shown cristae elimination [29,30]. However, while TEM can reveal precise intercellular compartment structures, it cannot capture dynamic processes in living cells. Using STED nanoscopy, we tracked changes in cristae density over time, which allowed us to see a sustained change caused by PLI in both cell types. After 24 hours of PLI-inducing cytokine exposure, PL-KCs showed a clear decrease in cristae density, while PL-FCs exhibited early signs of this process. Cristae restructuring can release mtDNA, activating cGAS/STING and producing type I interferons [31]. Increased IFN-α and IFN-β production was also found in the media of PL-cells. These interferons, crucial in psoriasis initiation, are overexpressed in patients’ blood and lesional skin [32]. Psoriatic patients also show high blood mtDNA levels [33]. These findings suggest a possible link between the disorganisation of cristae and psoriatic inflammation, though the activation of the mtDNA/cGAS/STING/IFN pathway in PL-KCs and PL-FCs is not yet clarified.

In our study, the reduced mitochondrial area and cristae density correlated with decreased mitochondrial respiration in PL-KCs. This is in line with the observed impact of cristae reorganisation on respiratory chain complex assembly and, consequently, mitochondrial efficiency [34,35]. However, this correlation is unclear in PL-FCs, where cristae density reduction did not affect respiration after 24 hours, possibly due to delayed response of this process. Interestingly, in PL-KCs, decreased mitochondrial respiration is accompanied by increased Δψ_m_ after 24 hours. This indicates a reverse electron transport (RET), potentially driven by ATPase inhibition. Our data shows decreased ATP synthesis in PL-KCs, which may minimise the utilisation of the Δψ_m_ across the inner mitochondrial membrane, therefore contributing to its elevation [36]. Similar alterations were seen in macrophages under bacterial inflammation [37]. Moreover, we also showed a correlation between MitoROS and respiratory chain activity. In PL-FCs, the progressive rise of MitoROS resulted in mitochondrial respiration decrease. Other researchers reported similar changes in dendritic cells treated with imiquimod to induce psoriasis-like inflammation [8].

PL-KCs exhibited early glycolysis dysfunction, while PL-FCs showed late glycolysis activation. Previous study reported that miR-31, which is overexpressed in psoriatic epidermis, changes metabolic enzyme expression, resulting in reduced glycolysis [38]. PLI-inducing cytokines can trigger miR-31 expression via NF-κB in HaCaT cells [39], suggesting a similar mechanism in PL-KCs. Other researchers have found that psoriatic skin shows increased glycolysis [40]. Understanding these mitochondrial response patterns in KCs and FCs may help identify new targets for psoriasis treatment.

PLI significantly increased psoriasis-related inflammatory mediators in both cell types. Specifically, PL-KCs had higher Elafin and IL-1β concentrations, aligning with their primary expression in psoriatic epidermis [41,42]. In turn, PL-FCs showed higher concentrations of IL-6 and IL-8, which is linked to recruiting IL-17A-releasing cells [43]. Despite variations in serum inflammatory mediator levels in psoriasis patients, our results match *in vivo* data ranges [44,45]. Cell-type specific mitochondrial responses to PLI indicate higher sensitivity of KCs and the delayed reaction of FCs, likely due to their distinct roles. KCs, forming a protective barrier, respond rapidly to stimuli, while FCs, maintaining the extracellular matrix, have slower signalling and metabolic demands [46].

In conclusion, our study reveals that psoriasis-like inflammation in keratinocytes and fibroblasts induces mitochondrial changes, such as mitochondrial respiration inhibition, increased mitochondrial membrane potential and MitoROS production, mitochondrial fragmentation, swelling, and cristae remodelling (Fig. 8). These changes may contribute to regulating proinflammatory mediator production, potentially revealing new targets for managing psoriasis. Further investigations are required to confirm the mechanistic link between mitochondrial changes and inflammatory responses, as well as the specific role of mitochondrial cristae in modulating mitochondrial functions. Additionally, employing gene-specific modulation strategies and primary cell models will be essential to extend the current findings.

**Figure 8.**
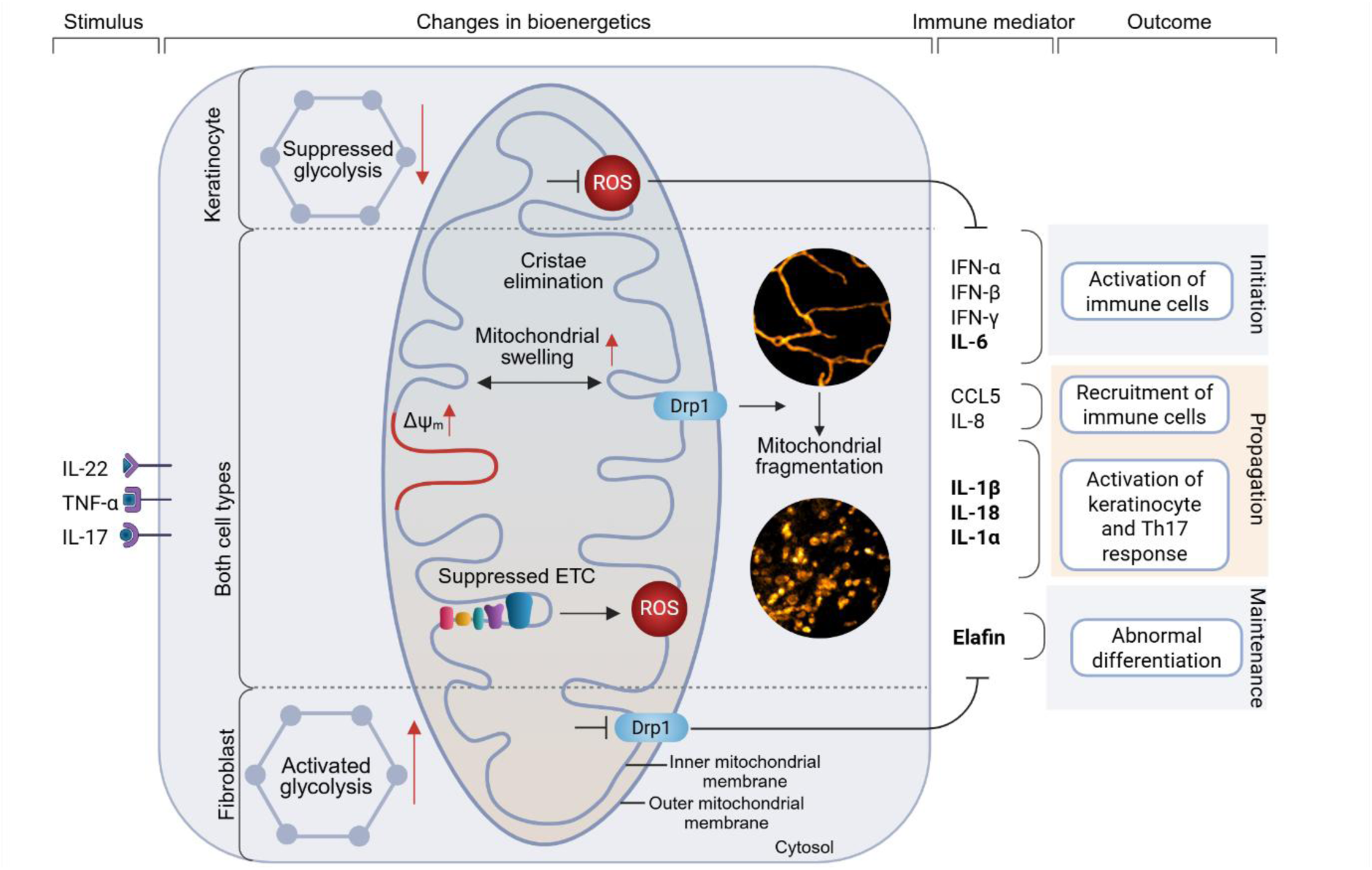
Mitochondrial changes in psoriatic-like keratinocytes and fibroblasts. Stimulation with IL-22, TNF-α and IL-17 induces secretion of proinflammatory mediators crucial for psoriasis initiation, propagation and maintenance. Under these conditions, both cell types exhibit increased mitochondrial swelling, cristae elimination, increased mitochondrial membrane potential (Δψ_m_) and suppressed mitochondrial electron transport chain (ETC). Also, this condition activates mitochondrial reactive oxygen species (ROS) production and Drp1-associated mitochondrial fragmentation, contributing to the secretion of IL-6, IL-1β, IL-18, IL-1α and Elafin. The cytokine induction pathway differs between the two cell types: in keratinocytes, it is mediated by mitochondrial ROS, while in fibroblasts, it is related to Drp1 activity. Also, psoriasis-like inflammation suppresses glycolysis efficiency in keratinocytes but activates it in fibroblasts. Illustrations were created using BioRender.com.

## Materials and Methods

The names and catalogue numbers of all reagent manufacturers are provided in Table 1.

**Table 1.**
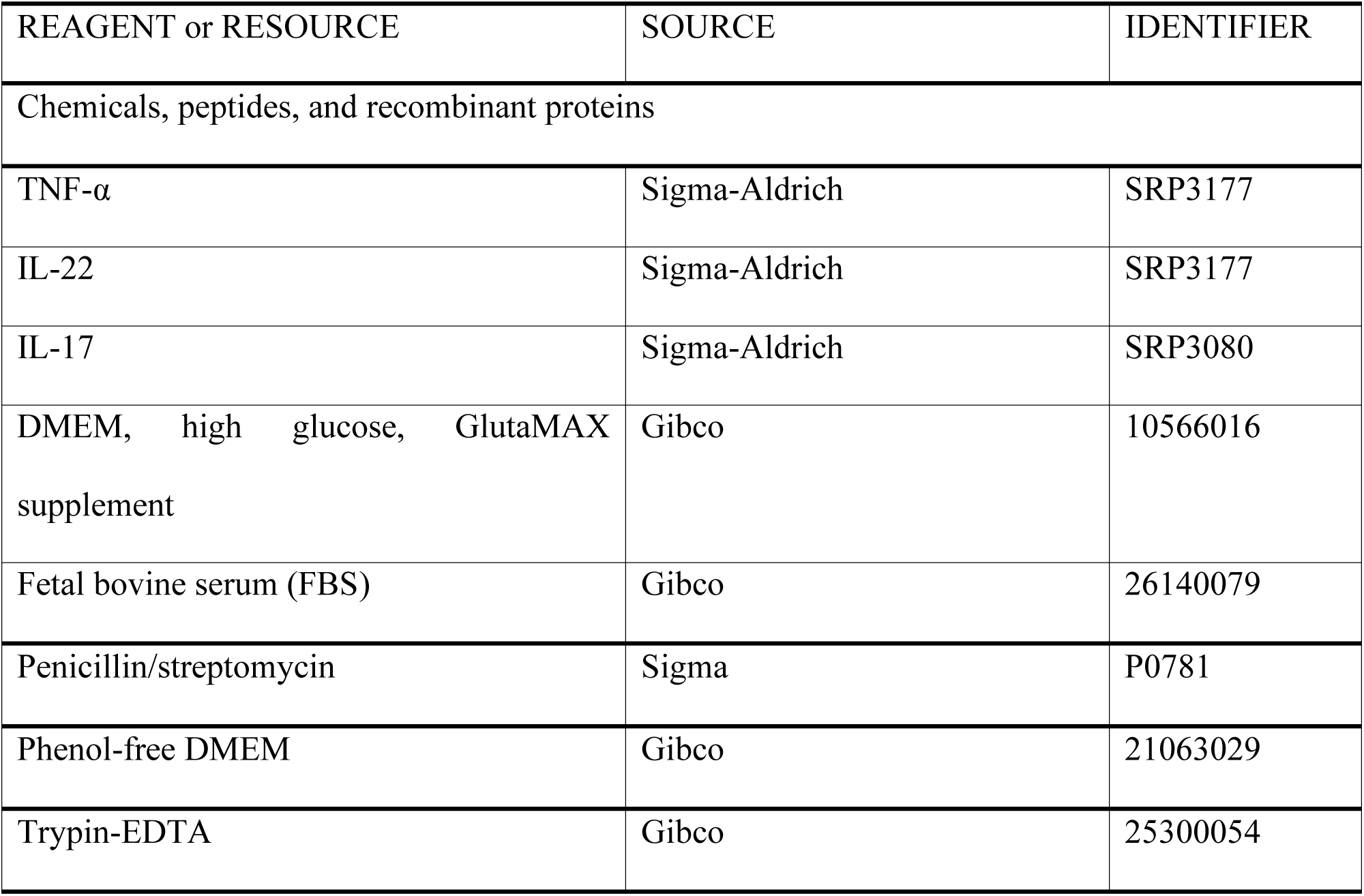
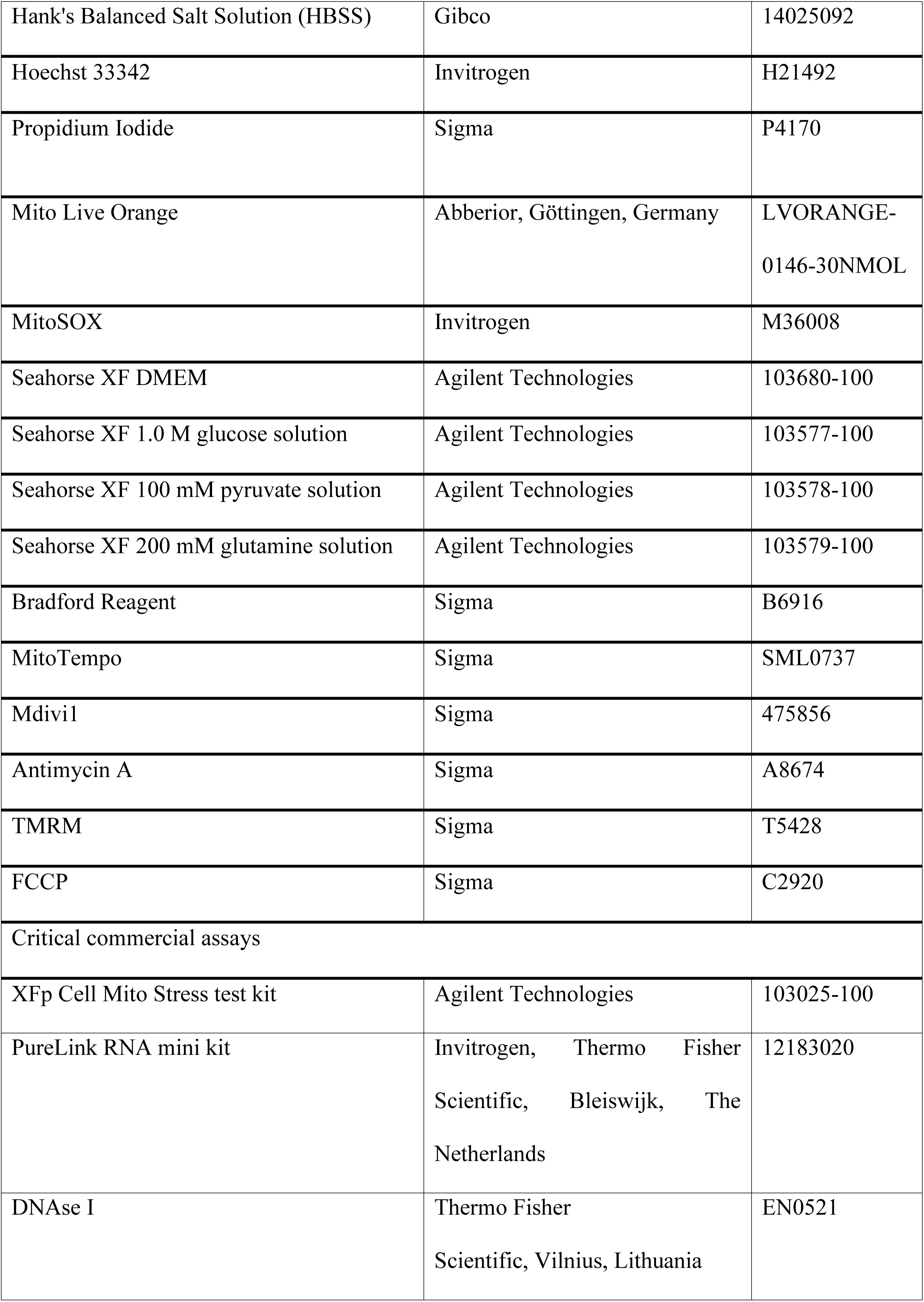
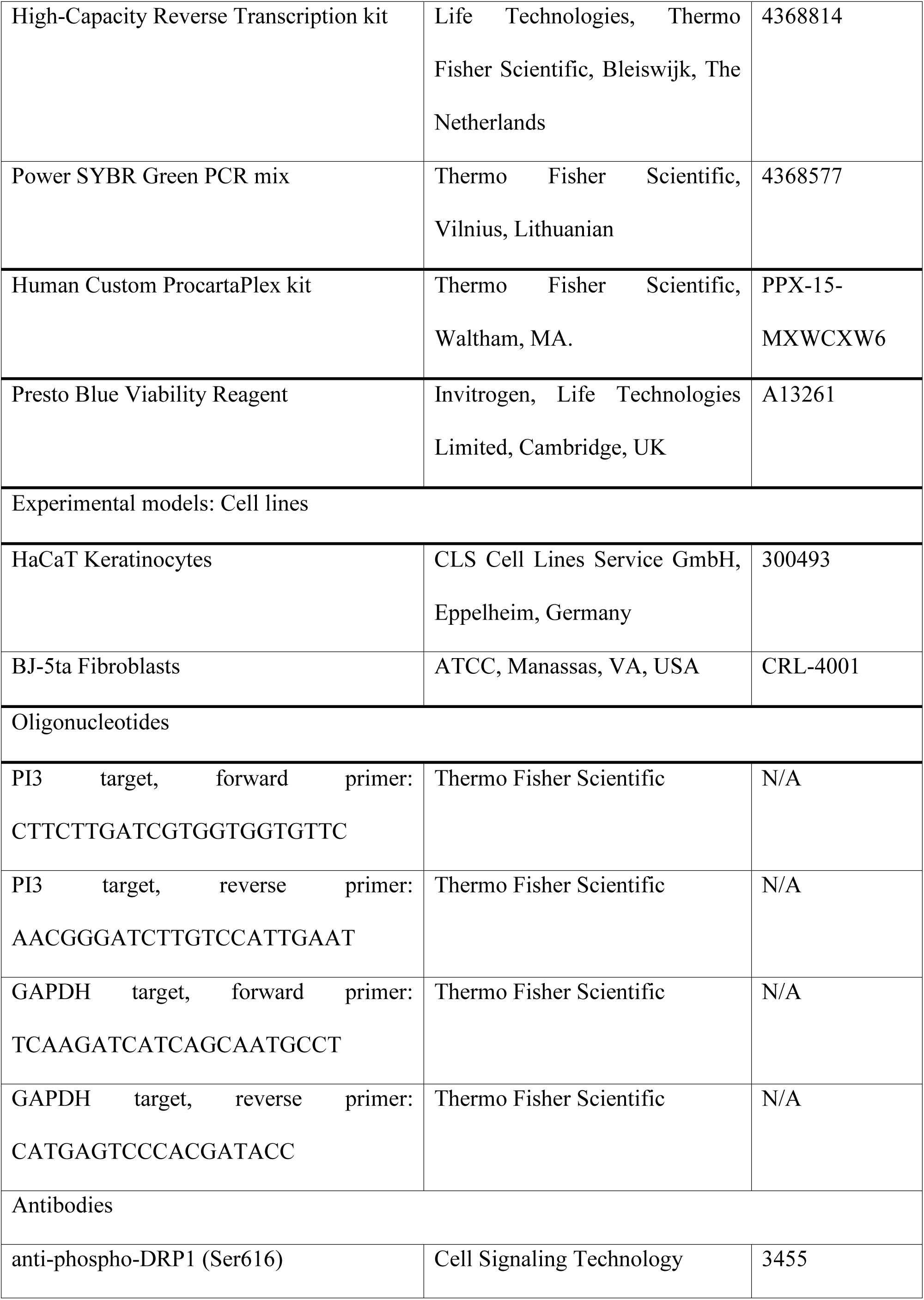
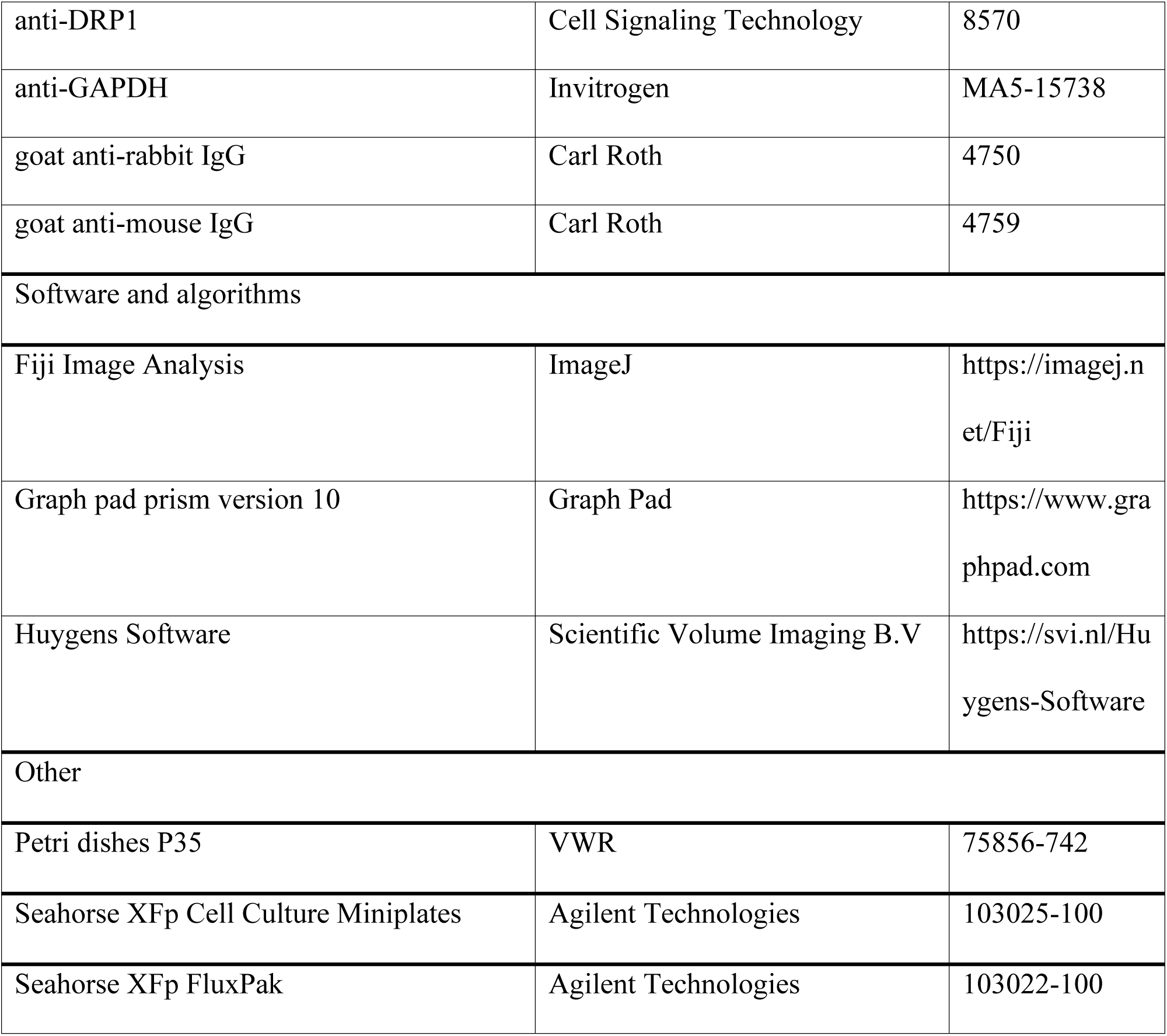
Key resources.

### Cell cultures and treatments

Human skin keratinocytes (HaCaT, RRID:CVCL_0038, CLS Cell Lines Service GmbH, Eppelheim, Germany) and fibroblasts (BJ-5ta, RRID:CVCL_6573, ATCC, ATCC, Manassas, VA, USA) were maintained in complete culture media containing DMEM with 10% fetal bovine serum and 1% penicillin/streptomycin at 37 °C. For PLI induction, the next day after plating, cell media was supplemented with cytokines for 24, 48 of 72 hours: TNF-α (5 ng/ml), IL-22 (10 ng/ml), IL-17 (10 ng/ml), from Sigma-Aldrich. PBS was used as a vehicle control. Other treatments: MitoTempo, a MitoROS scavenger, was used at 20 µM, added 1h before cytokine treatment, and co-incubated for 24 or 72 hours. Mdivi1, a mitochondrial division inhibitor, was used at 25 µM and was added 24 hours before the experiment. Both cell lines were confirmed to be mycoplasma-free through regular monitoring throughout the experiments.

### Gene expression

Total RNA was isolated using the PureLink RNA mini kit, followed by DNAse I treatment to remove DNA traces. Reverse transcription was performed using the High-Capacity Reverse Transcription Kit. Real-time PCR was conducted with Power SYBR Green PCR mix on a Rotor-Gene-Q System (Qiagen) using the 2^(-ΔΔCt)^ method for relative gene expression analysis, with glyceraldehyde 3-phosphate dehydrogenase (GAPDH) as the endogenous control. Results are depicted in the Log2 Fold Change form.

### Protein secretion

For protein secretion analysis Human Custom ProcartaPlex kit was used with Luminex 200 instrument (Luminex Corp, Austin, TX) following the manufacturer’s instructions. xPONENT software facilitated assessment. Values were presented as mean fluorescent intensities (MFIs) or concentrations (pg/ml) using a standard range equation for the relevant analyte. Values exceeding the detection limit were substituted with the highest standard concentration.

### Cellular respiration

Cells were seeded in Seahorse XFp cell culture plates at 5000 cells/well and, the next day were treated with PLI-inducing cytokines for 24h. For 72-hour treatment, cells were plated at 1700 cells/well. The bioenergetic function was evaluated using the Seahorse XFp Analyser (Agilent Technologies, Santa Clara, CA, USA) with the Seahorse XFp Cell Mito Stress Test Kit. The assay was conducted according to the manufacturer’s instructions. Briefly, following treatment incubation, cells were washed with XF DMEM (10mM glucose, 1 mM pyruvate and 2 mM glutamine) and incubated for 1 hour in a non-CO_2_ incubator. Final concentrations of electron transport chain-targeting compounds of the MitoStress test kit are as follows: 1.5 µM oligomycin, 2 µM for FCs, 1 µM for KCs FCCP, and 0.5 µM for rotenone and antimycin A. Results were normalised to total cellular protein content determined with the Bradford reagent (Sigma) and assessed using a multifunctional plate reader Infinite 200 Pro Nano Plex (Tecan, Männedorf, Switzerland).

### Cell metabolic activity and viability assessment

Cells were seeded at 5000 cells per well in 96-well plates and treated as previously described. Metabolic activity was assessed using PrestoBlue^TM^ Cell Viability Reagent following the manufacturer’s instructions. After 24, 48, and 72 hours of incubation, cell media was replaced with a solution containing 10% PrestoBlue reagent and 90% cell culture media. After a 40-minute incubation at 37 °C in the dark, fluorescence was measured using a plate reader (Infinite 200 Pro M Nano Plex, Tecan, Männedorf, Switzerland) at excitation and emission wavelengths of 560 nm and 590 nm, respectively. For cell viability and number assessment, cells were stained with nuclei dyes Hoechst33342 and propidium iodide (PI). Cells stained only with Hoechst33342 (blue nuclei) were considered viable, while cells with both blue and red (PI) nuclei classified as necrotic. Cells were incubated with 4 μg/mL Hoechst33342 and 7 μM PI for 15 min, followed by one wash with fresh media. They were analysed with fluorescent microscopy (Olympus APX100) and the total number of cell was counted in 5 randomly chosen regions with Fiji software.Viability was expressed as ratio of live and dead cells in a popululation. Control group was standardised to 100%.

### Mitochondrial superoxide and membrane potential determination by fluorescent microscopy

For MitoROS assessment, cells were stained with 2 μM MitoSox Red in HBSS at 37 °C for 30min, followed by two washes. Antimycin A, a superoxide inducer, was used as a positive control at 50 µM for 10min.

For Δψ_m_ determination, cells were incubated with 15 nM TMRM in phenol red-free DMEM at 37 °C for 25min. FCCP, a protonophore that disrupts mitochondrial membrane potential, was used at a concentration of 10 µM for 25min, concurrently with the TMRM stain, serving as a positive control. Imaging is performed without washing the dye (non-quenching/redistribution mode), allowing to estimate steady state Δψ_m_ [47].

For both assays fluorescence intensity was analysed using an Olympus APX100 microscope and analysis was conducted using Fiji software, where threshold-based segmentation (Huang algorithm) was applied. Control group was standardised to 100%.

### Mitochondrial superoxide determination by flow cytometry

Cells were grown in 24-well plates and treated with PLI-inducing cytokines for 24 and 72 hours. 200,000 cells per sample were collected and stained with 2µM in HBSS at 37 °C for 20min. Staining procedure was conducted according to the protocol [48]. Corresponding unstained cells were used as negative controls to define the background fluorescence. Flow cytometry files were processed using FlowJo v10 software (Becton, Dickinson and Company, Franklin Lakes, NJ, USA). The fluorescence intensities were registered for at least 10,000 cells per sample. Values of Geometric Mean of MitoSox Red were used to quantify ROS content in each sample. Signals were normalised by subtracting the Geometric Mean of the MitoSox Red value of unstained samples from the values of stained samples. Flow cytometry analysis was performed with a BD FACSMelody cell cytometer using BD FACSChorus software (version 3.0, Becton, Dickinson and Company, Franklin Lakes, NJ, USA).

### Mitochondrial morphometry

Cells cultured in glass-bottom Petri dishes at a density of 40,000. To visualise mitochondrial structures, cells were stained with 100 nM Mito Live Orange for 40min in a medium without FBS, followed by washing and imaging in HBSS using Zeiss Axio Observer.Z1 (Zeiss, Oberkochen, Germany) and Olympus IX83 (Olympus Corporation, Tokyo, Japan) fluorescent microscopes. FCCP at 0.2 µM for 10min induced extensive fragmentation. Mitochondrial morphology parameters (Figure 2a) were quantitatively assessed using the Fiji plugin Mitochondrial Network Analysis (MiNa) toolset [49], including mitochondrial area, mean of network branches, mean branch length, and mean of summed branch lengths. At least 10 cells per experimental group in three repeats were analysed.

### Cristae imaging and analysis

Cells were plated and stained with Mito Live Orange, as indicated previously. Imaging was performed using a STEDYCON system (Abberior Instruments GmbH, Göttingen, Germany) equipped with an Olympus IX83 microscope, a 100x oil immersion objective with a NA 1.45. Cristae area, density and spacing were assessed following steps depicted in Figure 3a). For cristae imaging, image acquisition employed 10-30 pixel sizes. Image size varied from 5 µm x 5 µm to 10 µm x 10 µm. Fluorescence signal accumulation employed 2-5 line steps with dwell times of 10 µs, and a pinhole set at 40 µm. For time-lapse recording, images were captured with a frame interval of 10-15 s for 60s. Compensation for photobleaching was carried out by bleach correction feature in Fiji. Subsequently, acquired images were subjected to deconvolution using Huygens software (Scientific Volume Imaging B.V., Hilversum, The Netherlands).

For average crista area measurement, cristae were segmented using the Trainable Weka Segmentation Tool in Fiji, following the guidelines provided in a paper [50]. Briefly, the plugin generated a probability map highlighting potential cristae areas compared to the background. These areas were then thresholded to create binary masks and processed using the “Watershed” function. Afterward, the “Analyse Particles” function outlined and marked cristae structures as regions of interest (ROIs). Finally, these structures were overlaid onto the original image and measured. Cristae spacing and density were assessed by analysing fluorescence intensity profiles along 1-2 µm line segments. For mitochondrial thickness assessment, intensity profiles were plotted across the mitochondrial width. At least three line profiles were measured for cristae analysis, while another three were used to evaluate the width of the mitochondrial filament in each image. ROIs for plotting fluorescence intensity profiles depended on cristae clarity, where mitochondrial filaments were not entangled, coiled, or of the focal point. Intercristae distance was determined by the distance between fluorescence peaks, and cristae density was calculated as the number of peaks per 1 µm segment. Fluorescence peaks were identified using Fiji’s BAR macro “Find peaks”. Time-dependent analysis involved fluorescence intensity profile measurement in each frame of the time-lapse. At least 10 cells per experimental group in four experimental repeats were examined, and analysis was conducted on randomly chosen regions. For time-lapse analysis, 4-6 video recordings were analysed.

### Drp1 detection using Western Blot

Cells were seeded at 70.000 cells per well in 24-well plates and treated for 24 hours with PLI-inducing cytokines. Cells were lysed, and whole-cell extracts were quantified by a Bradford Assay. For Western Blot, denatured protein samples (5 µg per lane) were prepared by adding 4× Laemmli sample buffer supplemented with 4% β-mercaptoethanol, followed by heating at 95 °C for 5 minutes. Proteins were separated by SDS–polyacrylamide gel electrophoresis (SDS–PAGE) and subsequently transferred onto nitrocellulose membranes (0.2 µm pore size) using a wet transfer system. After transfer, membranes were rinsed thoroughly with TBST (Tris-buffered saline containing 0.1% Tween-20), then blocked in 3% BSA in TBST for 1 hour at room temperature (RT) with gentle agitation (30–50 rpm).

Membranes were incubated overnight at 4 °C with primary antibodies diluted in blocking buffer: anti-phospho-DRP1 (Ser616) (Cell Signaling Technology, Cat# 3455, 1:1000), anti-DRP1 (Cell Signaling Technology, Cat# 8570, 1:1000), or anti-GAPDH (Invitrogen, Cat# MA5-15738, 1:4000). The following day, membranes were washed three times with TBST (5 minutes each), then incubated for 1 hour at RT with horseradish peroxidase (HRP)-conjugated secondary antibodies diluted 1:10,000 in blocking buffer: goat anti-rabbit IgG (Carl Roth, Cat# 4750) or goat anti-mouse IgG (Carl Roth, Cat# 4759). After secondary antibody incubation, membranes were washed again in TBST to remove excess antibody.

Protein bands were visualised using enhanced chemiluminescence (ECL) detection reagents (Pierce™, Thermo Scientific) and imaged with the Alliance Q9 Advanced imaging system. Densitometric analysis was performed using NineAlliance™ software.

### Quantification and statistical analysis

Statistical analysis comprised at least three independent biological experiments with a minimum of two technical replicates each. The dots on each bar represent the average value from a single biological experiment. Pairwise comparisons used an unpaired, two-tailed Student’s t-test. For comparisons involving more than two groups, we conducted a one-way ANOVA with Fisher’s LSD test for inter-group comparisons, and Dunnett’s test for comparisons with the control group. Grouped data underwent analysis using two-way ANOVA, followed by Dunnett’s test. The data analysis utilised GraphPad Prism software, version 10.0 (GraphPad Software, Inc., USA).

## Author Contributions

Conceptualisation, A.J., and R.M; Methodology, G.K.; Formal analysis, G.K., M.U., S.H., M.K., J.Š., V.Š. and M.I; Investigation, G.K., M.U., S.H., M.K., J.Š., V.Š., R.I, J.S. and M, I.; Resources, A.J., R.M.; Writing - original draft preparation, G.K. and A.J.; Writing - review and editing, G.K., A.J. and R.M.; Visualization, G.K., and A.J.; Supervision, A.J., R.M.; Project Administration: A.J.; Funding acquisition: A.J. All authors have read and agreed to the published version of the manuscript.

## Acknowledgements

This work was supported by the Research Fund of the Lithuanian University of Health Sciences. This project has received funding from the Research Council of Lithuania (LMTLT), agreement No: S-A-UEI-23-7.

## Data Availability

No datasets were generated or analysed during the current study. Any additional information required to reanalyse the data reported in this work paper is available from the lead contact upon request.

## Abbreviations

Drp1: Dynamin-1-like protein
ECAR: extracellular acidification rate
FCCP: Carbonyl cyanide-p-trifluoromethoxyphenylhydrazone
FCs: fibroblasts
IL: interleukin
KCs: keratinocytes
Mdivi1: mitochondrial division inhibitor
mitoROS: mitochondrial reactive oxygen species
OCR: oxygen consumption rate
PLI: psoriasis-like inflammation
PL: psoriatic-like
TNF-α: tumour necrosis factor α
TMRM: Tetramethylrhodamine methyl ester.

## Notes

**Conflict of Interest:** The authors declare no conflicts of interest.

### Competing Interest Statement

The authors have declared no competing interest.

### Summary of Updates

All figures revised; additional experiments performed; authors' list updated. This version has been peer reviewed and accepted for publication in FEBS journal

